# Hymenoptera associated eukaryotic virome lacks host specificity

**DOI:** 10.1101/2020.09.15.298042

**Authors:** Ward Deboutte, Leen Beller, Claude Kwe Yinda, Chenyan Shi, Lena Smets, Bert Vanmechelen, Nadia Conceição-Neto, Kai Dallmeier, Piet Maes, Dirk C de Graaf, Jelle Matthijnssens

**Affiliations:** KU Leuven – University of Leuven, Department of Microbiology, Immunology and Transplantation, Rega Institute for Medical Research, Division of Clinical and Epidemiological Virology, 3000, Leuven, Belgium; NIAID/NIH, Rocky Mountain Laboratories, Laboratory of Virology, Virus Ecology Unit, 59840, Montana, USA; KU Leuven – University of Leuven, Department of Microbiology, Immunology and Transplantation, Rega Institute for Medical Research, Laboratory of Virology and Chemotherapy, 3000 Leuven, Belgium; UGent – Ghent University, Department of Biochemistry and Microbiology, Laboratory of Molecular Entomology and Bee Pathology (L-MEB), 9000, Ghent, Belgium

## Abstract

Recent advancements in sequencing technologies and metagenomic studies have increased the knowledge of the virosphere associated with honey bees tremendously. In this study, viral-like particle enrichment and deep sequencing was deployed to detect viral communities in managed Belgian honey bees. A substantial number of previously undescribed divergent virus genomes was detected, including a rhabdovirus and a recombinant virus possessing a divergent *Lake Sinai Virus* capsid and a Hepe-like polymerase. Furthermore, screening > 5,000 public sequencing datasets for the retrieved set of viral genomes revealed an additional plethora of undetected, divergent viruses present in a wide range of Hymenoptera species. The unexpected high number of shared viral genomes within the Apidae family and across different families within the order Hymenoptera suggests that many of these viruses are highly promiscuous, that virus sharing within and between Hymenoptera families occurs frequently, and that the concept of species-specific viral taxa inside the Hymenoptera should be revisited. In particular, this estimation implies that sharing of several viral species, thought to be specific for bees, across other eukaryotic taxa is rampant. This study provides important insights on the host taxonomical breadth of some of the known “bee viruses” and might have important implications on strategies to combat viruses that are relevant to pollinators.

## Introduction

The European honey bee (*Apis Mellifera*) forms a central hub in ecosystem maintenance, resilience and diversity. Aside from the economically valuable products, such as honey and nectar (1,2), managed bee colonies together with other insects contribute tremendously to pollination (3) and play a key role in global agricultural production (4). In the past decades, pressures on both managed and wild bees have increased vastly and there is evidence for declining trends in pollinator populations globally (5,6). These pressures encompass ecological factors such as habitat loss (7), pollution (8), pesticide use (9,10) and adverse agricultural practices (11), but biological factors including bacterial, parasitic, and viral infections (12–15), also play a pivotal role. Recently, more attention is being given to the microbiota and their influence on bee health, development and homeostasis (16–18), and it has been shown that the microbiota can be exploited to protect bees from other pathogens (19). The influence these factors have can be cumulative or even synergistic. For example, it has been shown that pesticide use can perturb the expression of essential immunocompetence genes, increasing the probability of microbial infections (20). Perhaps the best example for mutual synergistic factors detrimental for bee health confine parasitic and viral infections. The worldwide spread of the *Varroa destructor* parasite facilitated *Deformed wing* virus (DWV) infections by acting as an active vector (where the virus can replicate in both the vector and the host) (21). Parallel to its role as viral vector, it has been shown that the *V. destructor* parasite can also influence the immune status of its host (22). Globalization of *V. destructor* and concomitant DWV infections raised the question what influence DWV plays in colony health. Recent studies have revealed an association between DWV infections and colony health status (23–25). Despite the worldwide dominance of DWV, other RNA viruses have been shown to be highly virulent, resulting in a strong phenotype in infected bees. Acute bee paralysis virus (ABPV), Black queen cell virus (BQCV) and Sacbrood virus (SBV) are all members of the order *Picornavirales* that have a detrimental effect on colony health once they infect a hive (26). Scattered information suggests that some of these viruses are not to be restricted to honey bees, but also infect and replicate in other members of the Apidae family. Spill-over events from managed honey bees into bumblebee species have been described for DWV, BQCV, ABPV, SBV and Lake Sinai viruses (LSV) (27–30), whereas honey bee viruses have also been described in ants (Formicidae) (31) and wasps (Vespidae) (32). Recent advancements in sequencing technologies and metagenomics have accelerated virus discovery in bees and a number of studies have attempted to describe the viral diversity associated with bees. These studies were able to expand the range of known honey bee viruses significantly and aside from numerous viruses belonging to the order *Picornavirales,* numerous other RNA viruses have been discovered belonging to the orders *Bunyavirales, Mononegavirales* (containing the family *Rhabdoviridae*) and *Articulavirales* (containing the family *Orthomyxoviridae*), and several unclassified RNA viruses such as LSV (33–37). DNA viruses have also been described, such as Apis mellifera Filamentous virus (AmFV) (38), and numerous single-stranded DNA viruses (39). While these sequencing efforts have vastly increased the number of known honey bee related viruses, the relevance of most of these viruses remains enigmatic. In this study, we first describe the eukaryotic viruses present in > 300 Belgian bee colonies collected in the framework of the EpiloBEE study (40) in 2012 and 2013. We place these results in the context of other known insect viruses. Finally, by screening more than 5,000 public RNA sequencing datasets, we shed light on the sharing of (bee) viruses between different members of the order Hymenoptera and within the Apidae lineage.

## Results

### Eukaryotic virus identification yields previously known and unknown honey bee viruses

Viral-like particle enrichment (41) and Illumina sequencing was performed on pooled samples derived from 300 weak and healthy (as defined by the EpiloBEE study (40)) managed honey bee colonies in Flanders, Belgium as described before (42). After sequencing and *de novo* assembly of the individual libraries, redundancy of the retrieved contigs was removed by collapsing sequences with 97% nucleotide identity over 80% of their length. Subsequently, the non-redundant contig set was annotated using DIAMOND (43) against NCBI’s NR database. Viruses were taxonomically classified using the lowest-common ancestor algorithm implemented in Kronatools (44). Sequences showing similarity to bacteriophages were omitted from this analysis. Genome coverage values were obtained by mapping the sequencing reads per sample back to the non-redundant contig set. Clustering analysis on the viral coverage matrix revealed a distinct clustering pattern between samples derived from weak and healthy colonies, although with a very small biological relevance (adonis test, R^2^ = 0.035, p-value = 0.0042) (Fig. 1A). The log-transformed coverage matrix showed that the vast majority of viral reads could be attributed to the family *Iflaviridae,* of which DWV is a member (Fig. 1B). The second most prevalent viral family was the family *Orthomyxoviridae*. Several families containing plant and fungal viruses, such as *Partitiviridae*, *Chrysoviridae*, and *Tymoviridae*, were also recovered. The clustering pattern of the coverage matrix reflected the adonis test results, showing most of the samples being dispersed by health status and although one healthy cluster containing mainly unclassified reads exists, the lack of monophyly implies no clear differences in composition with respect to the health status. In terms of absolute contig count, the most prevalent orders were (apart from unclassified sequences) *Picornavirales, Tymovirales* and *Mononegavirales* (supplemental fig. S1A) and the most prevalent families were (next to unclassified sequences) *Partitiviridae*, *Comoviridae,* and *Parvoviridae* (supplemental fig. S1B). There was no significant difference between the number of non-redundant contigs present in healthy and weak samples (Mann-Whitney U test, p-value = 0.32) (supplemental fig. S2). Only 30% of the non-redundant contigs had an amino acid similarity percentage with the best hit in the NR database higher than 90%, reflecting the divergent nature of the retrieved sequences (supplemental fig. S3). Species accumulation curves revealed a near horizontal asymptote, implying that viral sequence space was probed sufficiently (supplemental fig. S4). The relatively short length of the majority of retrieved viral sequences hampered a complete phylogenetic analysis (supplemental fig. S3). Therefore, an all-by-all TBlastX search was conducted using the retrieved non-redundant contig set complemented with a filtered viral Refseq set (see methods) as both query and bait. The resulting blast output was converted into a network using sequences as vertices, and hits as edges. A minimized-nested block network was constructed and visualized using the taxonomical information of the reference sequences (Fig. 1C). The vast majority of retrieved sequences clustered together in blocks with the order *Picornavirales*, although the orders *Bunyavirales*, *Mononegavirales* and *Tymovirales* were also represented substantially. Several contigs could not be assigned to any known order and represented unclassified (ds)RNA viruses or unclassified (circular) DNA viruses.

**Fig. 1.**
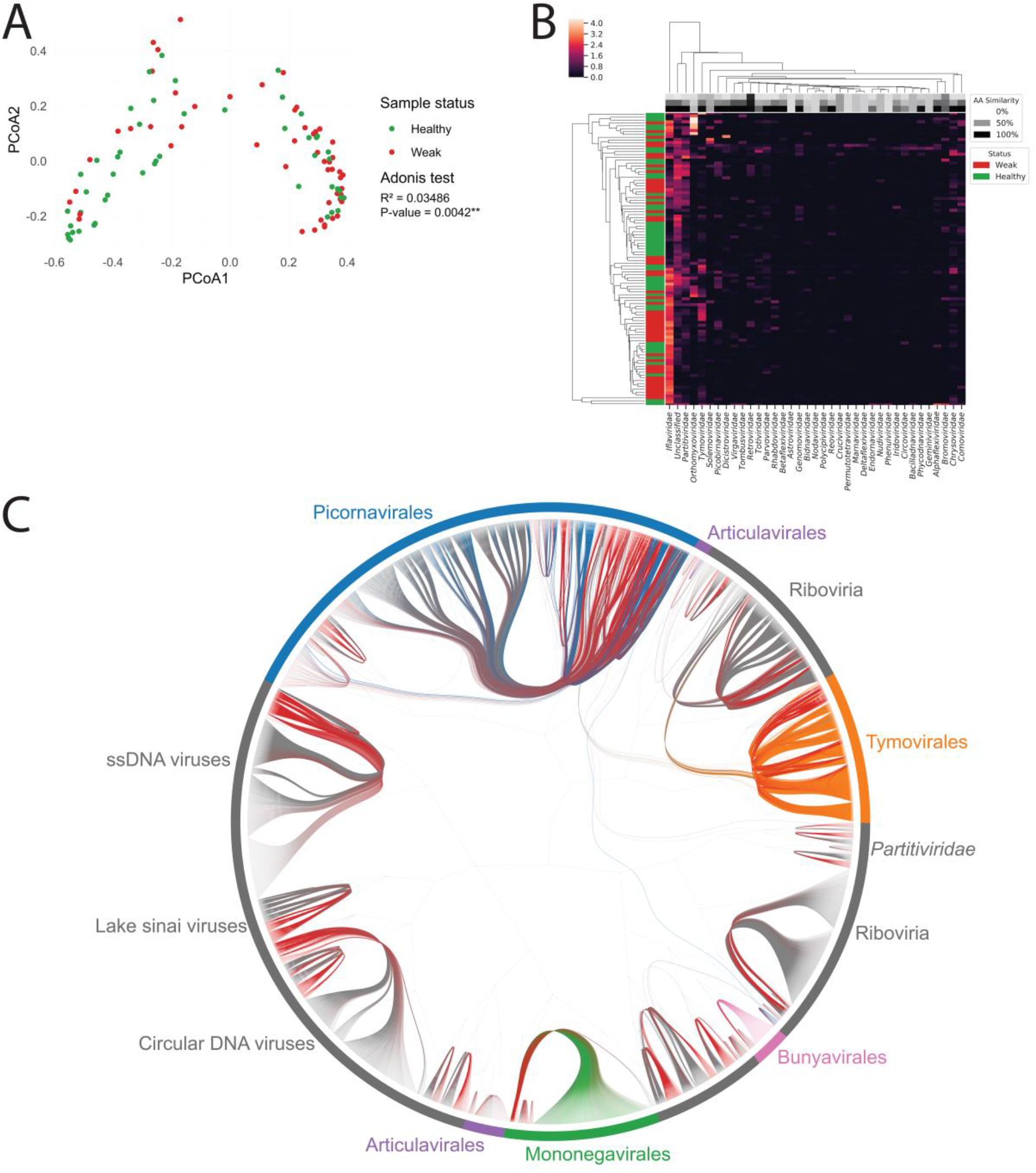
Belgian honey bees harbor a diverse range of known and novel viruses. (A) PCoA clustering using Bray-Curtis distances calculated on the viral coverage matrix derived from the Belgian samples (n = 102). Green dots reflect samples derived from healthy colonies; red dots reflect samples derived from weak colonies. The R squared and p-value obtained from the Adonis test are indicated on the right. (B) Average values per viral family of the log-transformed viral coverage matrix are depicted in a heatmap, clustered using Euclidian distances. The left column depicts samples derived from healthy (green) and weak (red) samples. The first three rows indicate the minimum (top), the average (middle) and the maximum (bottom) percentage of amino acid similarity of the contigs per viral family. (C) Minimized nested block network using retrieved sequences in this study (red) and known Refseq viruses (all other colors). Known orders are indicated in colors and unclassified reference sequences are indicated in gray.

### Phylogenetic analysis confirms the presence of known and divergent eukaryotic viruses in Belgium

To investigate the phylogenetic placement of a subset of the retrieved near-complete viral genomes, maximum clade credibility trees (MCC) were created using BEAST (45) (Fig. 2). Retrieved genomes from this study (orange tip labels, and listed in supplemental table 1) and from the short-read sequencing archive (SRA, NCBI) search (blue tip labels, see below), as well as reference sequences (green tips), were included based on sequence length and based on a BlastP search (see methods). Phylogenies were created for Rhabdo-like, Picorna-like, Bunya-like, Orthomyxo-like, Sinai-like, Partiti-like, Toti-like and Tymo-like viruses. One of the retrieved rhabdo-like viruses (Apis rhabdovirus1-Belgium) was nearly identical to the recently identified Apis rhabdovirus 1 (34), while the other Rhabdo-like virus (Apis rhabdovirus3-Belgium) has Diachasmimorpha longicaudata rhabdovirus as closest relative (but only had 38% amino acid identity for the L protein). The retrieved Picorna-like viruses reflect known bee pathogens clading in the families *Iflaviridae* and *Dicistroviridae,* such as DWV, SBV and ABPV, but also include more divergent sequences related to Nora-like viruses. A number of sequences clading together with plant infecting picornaviruses, such as several comoviruses were also retrieved. The retrieved Orthomyxo-like viruses are three closely related viruses (*Apis orthomyxovirus 1, 2* and *3-Belgium*), clustering together with other known thogotoviruses. These three viruses are nearly identical to the recently discovered Varroa Orthomyxovirus, with the exception of the nucleoprotein (35). Furthermore, five LSV-like viruses were retrieved, out of which four were very similar to other known Lake Sinai viruses (between 94% and 97% nucleotide similarity). Interestingly, the fifth identified *Lake Sinai virus* was initially identified as an Astro-like virus, but was shown to be a divergent recombinant virus with a ‘Hepe-like’ polymerase region (31% amino acid similarity with the non-structural protein of Culex Bastrovirus-like virus), and a Lake Sinai virus-like capsid (Lake Sinai virus, 35% amino acid similarity) (supplemental fig. S5). Sequence depth profiling indicated that this sequence was a true recombinant rather than an assembly artefact. The other retrieved (near-) complete viral genomes were most likely plant derived eukaryotic viruses, including Partiti-like viruses (24 sequences), Toti-like viruses (two sequences) and Tymo-like viruses (four sequences).

**Fig. 2.**
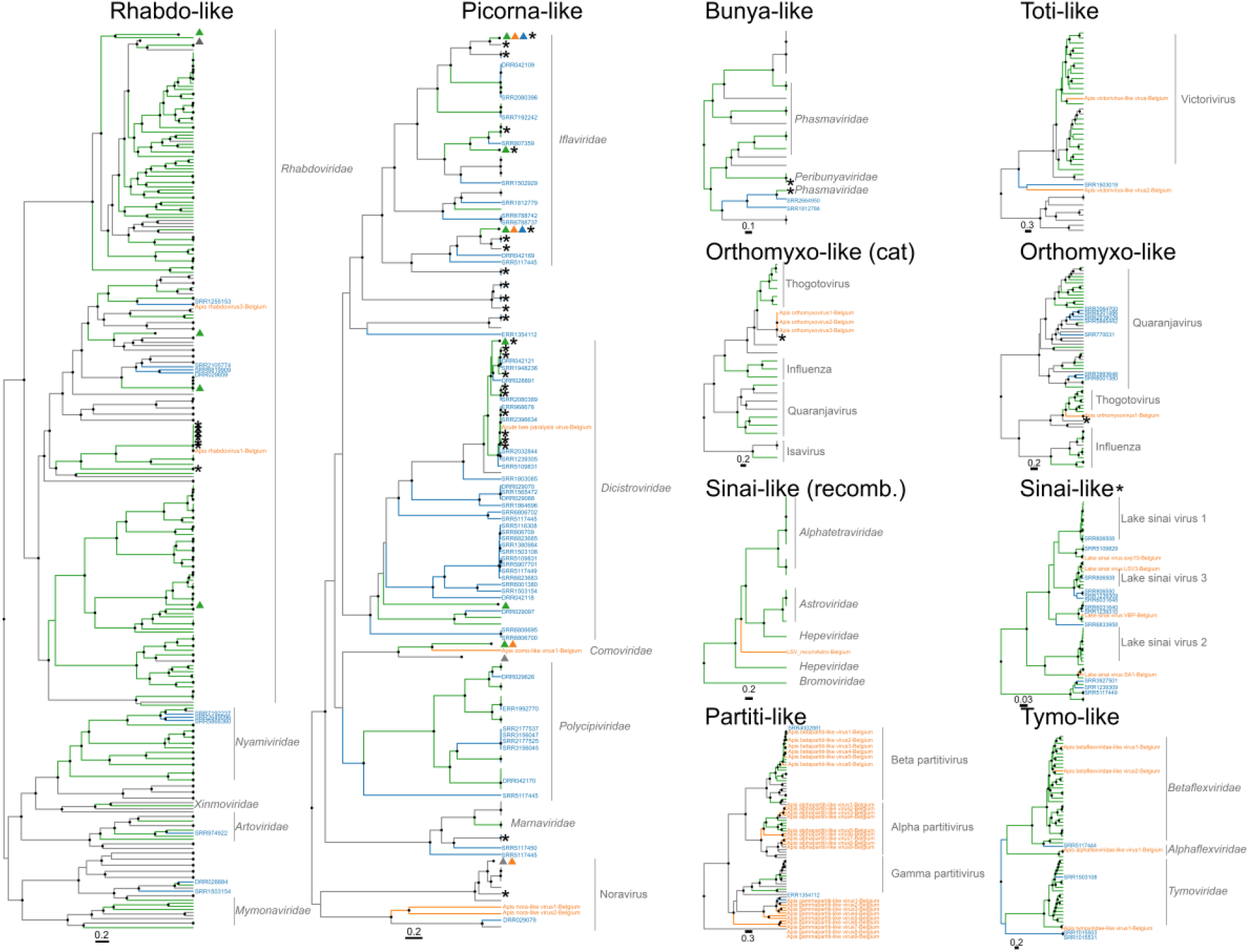
Phylogenetic analysis highlights the vast diversity of viruses identified in the Belgian samples. Maximum clade credibility trees for the best-represented groups of viruses retrieved in this study. Black circles on the nodes indicate posterior support values > 0.9. Viruses identified in the Belgian samples are indicated with orange tip labels, those identified through SRA searches are indicated with blue tip labels. Reference sequences that belong to a classified viral family or genus are indicated with green branches. Known honey bee viruses are indicated with an asterisk. One or more triangles indicate collapsed clades, and the colors are equivalent to the tip and branch colors. Clades that belong to the same family or genus are indicated with a gray line and in text. The ‘Rhabdo-like’ tree is built using the putative L protein. The ‘Picorna-like’ tree is built using the putative polyprotein (monocistronic viruses), the putative ORF1 (dicistronic viruses) or the putative replication polyprotein (Nora-like viruses). The ‘Bunya-like’ tree is built using the putative L protein. The ‘Toti-like’ tree and the ‘Partiti-like’ tree are built using the putative RdRP gene. The ‘Orthomyxo-like (cat)’ tree is built using a concatenated protein alignment of the putative PB2 – PB1 – PA – NP genes, while the ‘Orthomyxo-like’ tree is built using only the PB2 segment. The ‘Sinai-like (recomb.)’ tree is built using the putative polymerase gene of the astrovirus-LSV recombinant virus, while the ‘Sinai-like’ tree is built using the putative polymerase region of all the known LSV viruses (not including the recombinant). The ‘Tymo-like’ tree is built using the putative polyprotein gene.

### Re-screening of existing RNA sequencing datasets reveals untapped viral diversity within the Hymenoptera lineage

Since the recovered viral sequences included most of the known honey bee viral sequence space (Fig. 1C), the assumption was made that the non-redundant viral dataset we recovered was a good reflection of all known honey bee viruses. This dataset was used as bait to map a total of 5,246 RNA sequencing datasets found in the SRA database when using the query ‘Hymenoptera + RNA’. A dataset was considered to be ‘virus enriched’ when at least 100,000 reads mapped to the bait set. All datasets that met this criterium (1,331) were individually *de novo* assembled using SKESA (46) and viral sequences were identified and clustered as was described for the Belgian samples. An additional clustering step was performed, collapsing the non-redundant SRA-derived sequences together with the non-redundant Belgian sequence dataset. This resulted in the recovery of nearly 10,000 non-redundant putative viral contigs, out of which only 42.8% had an amino acid similarity with proteins in Genbank higher than 90% (supplemental fig. S6). Forward model selection analysis revealed that together, putative host taxonomy and location of the dataset could explain 33% of the variability observed within the coverage matrix (Fig. 3A). This result was further validated by the observation that hierarchical clustering on Euclidian distances revealed clusters of both eukaryotic host families and location within the coverage matrix (Fig. 3B). Viral taxonomy analysis revealed that the majority of the recovered viruses could be assigned to the orders *Picornavirales* and *Mononegavirales* (Fig. 3C). The retrieved viral contigs that fell below the abovementioned threshold of 90% amino acid similarity were included in the phylogenetic analysis and revealed ten previously undescribed Rhabdo-like viruses, and more than 50 previously undescribed Picorna-like viruses (Fig. 2, blue tip labels). Both these groups span multiple viral families. Another striking finding was the fact that seven previously undescribed PB2 segments of *Orthomyxoviridae-like* sequences were recovered (most closely related to the *Quaranjavirus* genus), indicating that this viral family is more strongly represented within the Hymenoptera lineage than was previously known. Furthermore, also Bunya-like, Toti-like, Sinai-like, Partiti-like and Tymo-like viruses were recovered (Fig. 2, blue tip labels).

**Fig. 3.**
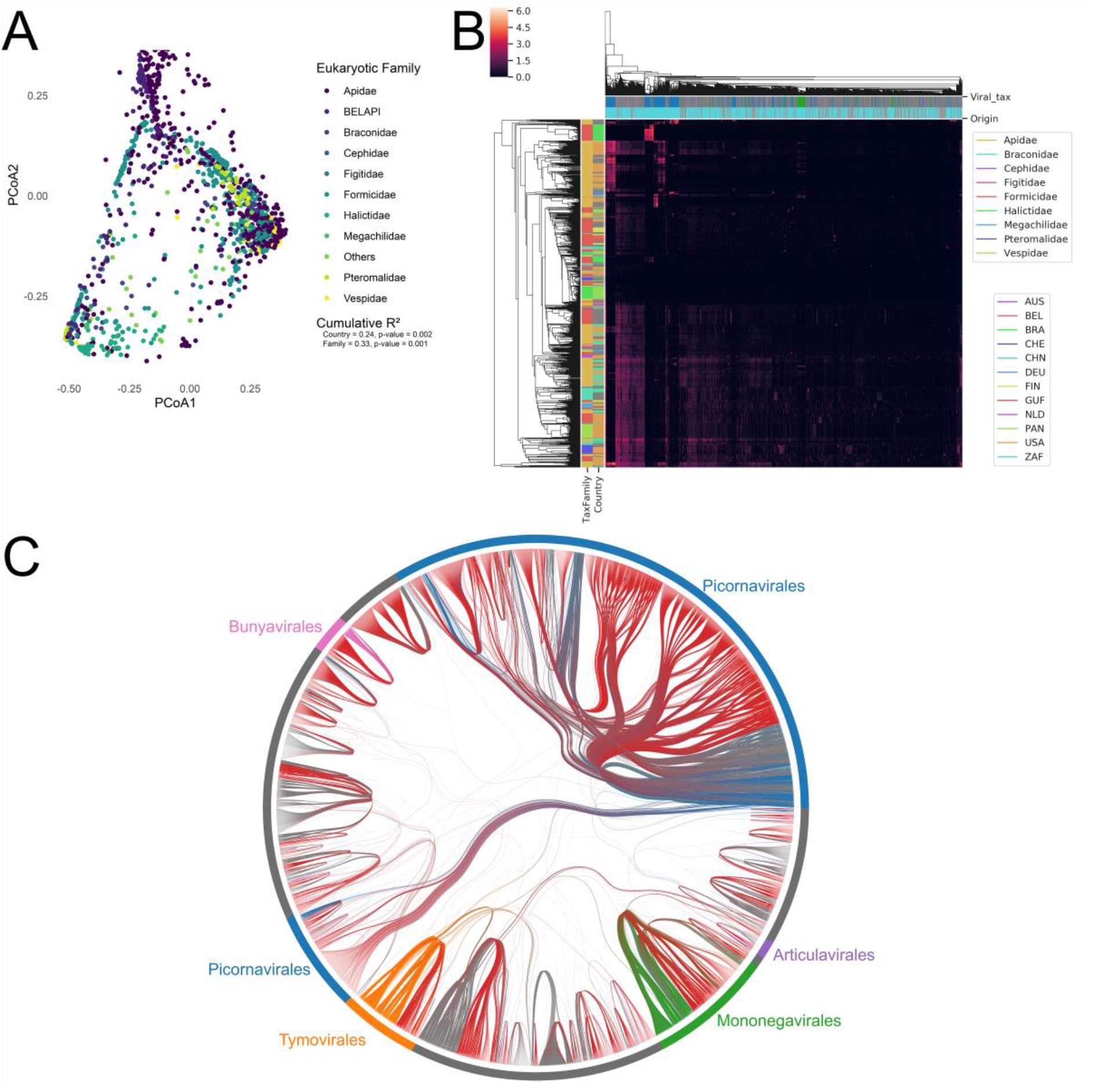
SRA searches shed light on the hymenoptera virosphere and reveal the wealth of undescribed viral sequences present in public datasets. (A) PCoA clustering using Bray-Curtis distances calculated on the viral coverage matrix derived from the Belgian samples clustered with the SRA screening results. Dots are colored per hymenoptera family. The cumulative R2 values reported are calculated by forward model selection using the OrdiR2step function after distance-based redundancy analysis. (B) Heatmap depiction of the log transformed viral coverage matrix, clustered using euclidian distances. Leftmost columns indicate the hymenoptera families and the geographical location of the samples. The top two rows indicate the viral taxonomical classification (with the same color per viral order as Fig. 3C) and the origin of the viral sequence (light blue indicates an SRA sample as origin, red indicates viral sequences found in the Belgian samples). (C) Minimized nested block network using the non-redundant sequences retrieved from the SRA searches (red) and known Refseq viruses (all other colors).

### Virus-sharing networks show a large number of virus sharing within the order of Hymenoptera

Although the discrimination between samples based on their location and eukaryotic taxonomy was significant (p-value 0.002 and 0.001, respectively) (Fig. 3A). The cumulative R2 value (0.33) indicates that a large majority of the variances within the datasets cannot be explained by aforementioned variables. This observation could imply that a large number of viruses are shared across hymenoptera families, that the variance within eukaryotic families in a specific country is large relative to the variance between these parameters, or a combination of both. To investigate the first possibility, the assumption was made that the host of a specific virus sequence was that of the sample of which the sequence cluster representative was derived. Hymenoptera families of which less than ten samples were obtained were grouped together into an ‘Others’ group and virus sharing was calculated in a pair-wise manner for all the possible combinations within the eukaryotic host families, and within the Apidae lineage. A substantial number of viruses were found to be present not only within the Apidae lineage but also shared over multiple eukaryotic host families (Fig. 4 A, B). Within the family Apidae, most viral sequences were shared between *Apis Mellifera* and *Lepidotrigona* species (1,050 viral sequences shared), between *Apis Mellifera* and *Apis florea* (938 sequences shared), and between *Apis Mellifera* and *Ceratina* species (729 sequences shared) (Fig. 4A). The majority of these shared sequences could be traced back to the order *Picornavirales,* with a total of 224 (21.3%), 497 (53.0%) and 130 Picorna-like sequences (17.8%) shared between these groups, respectively (Fig. 4C, blue edges). Beyond the family Apidae, substantial virus sharing was detected between the families Apidae and Pteromalidae (1,066 viral sequenced shared), the families Apidae and Cephidae (742 sequences shared), and the families Apidae and Braconidae (737 sequences shared). Concomitant with the situation between different Apidae species, the majority of shared viral sequences could be assigned to the order *Picornavirales,* with 201 (18.8%), 137 (18.4%) and 111 Picorna-like sequences (15.0%) shared between these groups, respectively (Fig. 4D, blue edges). Aside from Picorna-like sequences, evidence could also be found for sharing of viruses predicted to belong to the orders *Mononegavirales* (Fig. 4C,D, green edges) and *Tymovirales* (Fig. 4C,D, orange lines), although the number of shared viral sequences was on general an order of magnitude lower than those of the *Picornavirales* (39 Mononega-like viral sequences shared between Formicidae and Pteromalidae, and 27 Tymo-like viral sequences shared between *Apis Mellifera* and *Lepidotrogona).* Since a fraction of the recovered viruses are most likely infecting plants or reflect viruses not relevant for bees (Fig. 2), an additional analysis was ran with a number of the retrieved, near-complete, known bee viruses (AMFV, ABPV, BQCV, Kashmir Bee virus (KBV), DWV, LSV, *Apis Rhabdovirus* and *Apis Orthomyxovirus*), as well as the retrieved Nora-like viruses and other Orthomyxo-like viruses. Calculation of the fraction of positive samples revealed that most of the previously thought bee-specific viruses occur in multiple Apidae species but are also found within other Hymenopteran families (supplemental fig. S7). An attempt was made to quantify the host specificity of these viruses by calculating an Apidae specificity index (ASI), and an *Apis Mellifera* specificity index (AMSI) (Table 1). These indices revealed that some of the established bee viruses (ABPV, AMFV, BQCV and Quaranja-like orthomyxoviruses) show a low specificity for *Apis Mellifera* within the Apidae family (characterized by a low AMSI), and (with the exception of BQCV) were not restricted within the family Apidae (characterized by a low ASI). Other “established honey bee species” were shown to be highly specific for *Apis Mellifera,* and revealed a high AMSI (KBV, DWV and LSV). The recently discovered Apis rhabdoviruses and Nora-like viruses are found exclusively in *Apis Mellifera* within the family Apidae. The Apis rhabdoviruses are restricted within the family Apidae, but the retrieved Nora-like viruses are also highly prevalent in other Hymenoptera families (ASI 0.01, Table 1). Finally, the retrieved Quaranja orthomyxo-like viruses were highly prevalent in other Hymenoptera families and only to a limited extent in *Apis Mellifera* and the family Apidae (ASI of 0.04 and AMSI of 0.05, respectively). On the other hand, *Apis Orthomyxovirus 1* was slightly more honey bee specific, with an ASI and AMSI of 0.31 and 0.21, respectively.

**Fig. 4.**
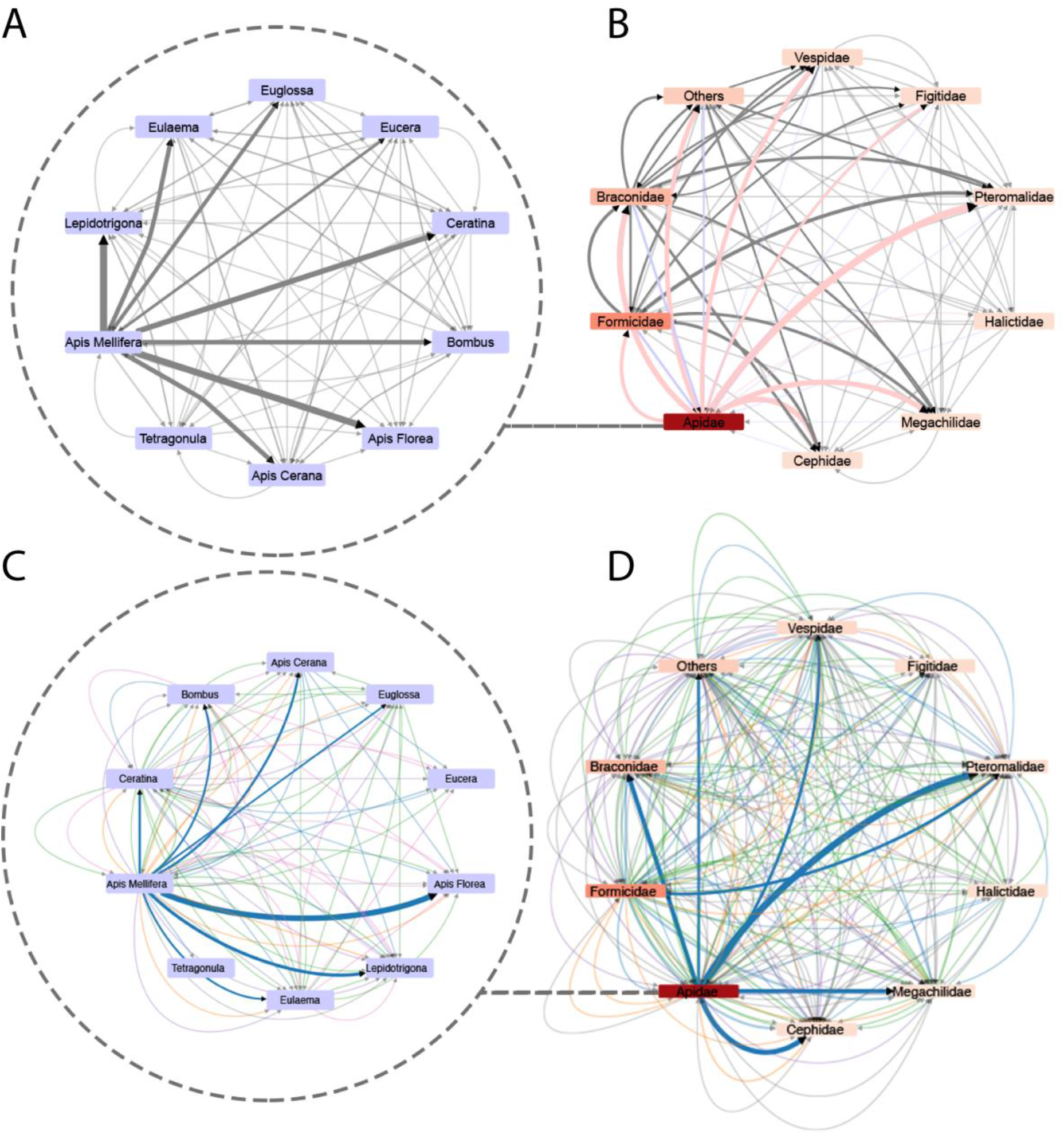
Cumulative viral sequence sharing network reflect the aspecificity of hymenoptera associated viruses. Networks reflecting the cumulative sharing of viral contigs between eukaryotic lineages. The networks inside the dashed circle (A,C) reflect viral sequence sharing within the family Apidae. The networks on the right (B,D) reflect sharing over different families within the order hymenoptera. Panels A and C and panels B and D both reflect the same networks, but both panels C and D reflect cumulative shared viral sequences broken up per assigned viral order (using the same color code as fig. 3C). Nodes in panels B and D are colored by number of representative virus contigs per eukaryotic lineage (ranging from 18 contigs (Cephidae) to 5,662 contigs (Apidae)). Edge thickness reflects the total shared contig count, ranging from 1 to 1,050 contigs (panel A), from 1 to 1,066 contigs (panel B), from 1 to 497 contigs (panel C), and from 1 to 201 contigs (panel D). Edge arrows indicate directionality, of which the root is the predicted host (the taxonomical group of which the virus sequence representative was derived from).

**Table 1.** Host (a)specificity of a selection of known bee viruses. The values reflecting how specific a known bee virus is for *Apis mellifera* (AMSI) and for Apidae (ASI). A value of 1 reflects complete lineage restriction. The number of viral contigs included per viral species is indicated with Contig number.

| Virus | Virus Abbreviation | Contig number | ASI | AMSI |
| --- | --- | --- | --- | --- |
| Acute Bee Paralysis virus | ABPV | 7 | 0.13 | 0.04 |
| Apis Mellifera Filamentous virus | AMFV | 8 | 0.06 | 0.05 |
| Kashmir Bee virus | KBV | 2 | 0.01 | 1.00 |
| Deformed Wing virus | DWV | 11 | 0.84 | 0.71 |
| Black Queen Cell virus | BQCV | 7 | 0.88 | 0.09 |
| Lake Sinai viruses | Sinaiviruses | 20 | 0.07 | 1.00 |
| Apis Rhabdovirus (1 and 2) | Rhabdo | 4 | 1.00 | 1.00 |
| Quaranja-like Orthomyxoviruses | Quaranja | 15 | 0.05 | 0.04 |
| Apis Orthomyxovirus 1 | Thogoto | 3 | 0.31 | 0.21 |
| Nora-like viruses | Nora | 3 | 0.01 | 1.00 |

## Discussion

This study, combined with other recent sequencing efforts, provides new insights into known and previously undescribed viruses associated with *Apis Mellifera.* Variance analysis revealed a significant, but biologically limited difference in the viral composition between weak and healthy colonies, and no significant difference in the total number of viral sequences derived from healthy and weak colonies could be detected. Genomes from a large number of viral families could be retrieved, of which a substantial part most likely includes plant viruses. While it cannot be excluded that some of the recovered divergent plant viruses constitute viruses actually infecting the bee, it is likely that the majority of these viruses reflect environmental contaminants. The host of the most closely related viral sequence can give an indication if these sequences are environmental contamination. The fact that numerous viral sequences belonging to families solely infecting plants were recovered in a large scale viral discovery study in insects indicates that this assumption does not necessarily hold true (47). The recent detection of viruses belonging to plant-specific viral families in mosquitoes reinforces this observation (48). Some of the retrieved viral sequences were very similar to recently discovered viruses (*Apis Rhabdovirus 1, Apis Orthomyxovirus 1*), increasing the likelihood that these are true honey bee viruses, and further confirming their presence in Belgium. Interestingly, a divergent recombinant Lake Sinai virus was found, comprised of a Hepe-like polymerase region, and a divergent Lake Sinai virus capsid. A novel divergent rhabdovirus (*Apis Rhabdovirus 3*) was also described. Additionally, full genomes for an orthomyxovirus (*Apis Orthomyxovirus 1*), very similar to a virus from a previous study from Levin *et al.* (35), was found in multiple individual libraries, and evidence for the presence of this virus was found in other Hymenoptera families (Table 1). Multiple sequence alignment-free network analysis implied that, despite the species accumulation curves reaching a plateau, many of the putative viral sequences retrieved were too fragmented to be included in a phylogenetic analysis (the number of sequences that made the threshold to be included in phylogenies was 188, while the network reflected 5,224 retrieved sequences). Furthermore, this analysis implies that the actual viral diversity exceeds what can be captured by regular phylogenetic analysis. Larger sample sizes and especially deeper sequencing efforts could help to fully elucidate the viral diversity associated with honey bees. Since the retrieved non-redundant viral sequence set encapsulates nearly all of the known and even more recently described viruses, this set was used to probe pre-existing Hymenopteran sequencing datasets for any bee-related viral signal. A total of 1,331 virus-rich RNA sequencing datasets were *de novo* assembled and screened for viruses. This approach revealed that these datasets harbor a substantial number of viruses that have been previously described (roughly 40%), but also that the amount of undescribed, divergent viruses is rampant. In concordance with the previous results, the viral sequences retrieved from the SRA search also suffer from fragmentation and incomplete sequencing. This observation is most likely the result of the fact that most RNA sequencing datasets included in the SRA search are transcriptome studies rather than metagenomic analyses, and that for most of them no wet-lab procedures for microbial or viral enrichment were performed. Despite this setback, multiple-sequence alignment free network analysis implied a massive hidden viral diversity within the Hymenoptera lineage (roughly 60% of the retrieved contigs were less than 90% similar to any other known virus in Genbank). Constrained ordination analysis showed that both the geographical origin and the taxonomical lineage of the host organism sequenced could explain a biologically relevant proportion (cumulative R2 = 0.33) of the variance within the viral coverage matrix. Since the included samples constitute a wide range of taxonomical host lineages, this proportion was below expectations and implies a substantial amount of viral sequences to be shared over eukaryotic Apidae species and Hymenoptera families. This hypothesis was confirmed by cumulative counting of the viral sequences over the different lineages included, based on a rather rigorous coverage threshold for presence/absence. This analysis revealed a non-trivial number of viral sequences, spanning all of the viral orders previously associated with honey bees, being shared across different lineages within the Apidae, but also over other families belonging to the Hymenoptera. Of all the sequences present in the total non-redundant viral dataset, 53% were shared with another taxonomical lineage (5139 shared sequences, 9655 in total). Since the included SRA dataset suffers strongly from sampling bias, this percentage is most likely an underestimation. Given this strikingly high number of virus sharing, the dataset was revisited with a subset of previously described honey bee specific viruses. Surprisingly, none of the tested viruses were lineage restricted to *Apis Mellifera*, with the exception of Apis rhabdovirus. Other viruses, such as Nora-like viruses, KBV and LSV were restricted to *Apis Mellifera* within the Apidae lineage (AMSI = 1.00) but were underrepresented relative to non-Apidae Hymenopteran families. The only viruses that were bee specific, *i.e.* having both a high AMSI and ASI, were DWV and Apis rhabdoviruses. These results imply that despite the recent sequencing efforts, many unknowns remain on viral diversity within the Hymenoptera lineage. Finally, the concept of “honey bee specific viruses” should be revisited, since most of the previously described viruses are not bee specific, neither are they restricted to the Apidae lineage.

## Methods

### Data and code availability

All relevant (intermediate) output files, metadata tables, fasta sequences, R code, Python code and jupyter notebooks are available on Github through the URL https://github.com/Matthijnssenslab/Bee_euvir. Intermediary output files too large to be hosted on Github are available through Zenodo (10.5281/zenodo.3979324). The raw sequencing data is available through the SRA database under project accession PRJNA579886. Accession numbers of the viral sequences included in the phylogenies will be made available in supplemental table S1. Accession information for the public datasets screened in this study are available in supplemental table S2.

### Sample preparation, pooling, VLP-sequencing and read processing

Samples were pooled and prepared for Illumina sequencing as described before, and the prokaryotic viruses in these pools were described previously (42). Briefly, samples were taken from the Flanders EpiloBEE study (40), from both sampling years (2012 and 2013), and 102 pools were constructed based on health status (defined retrospectively within the EpiloBEE study, with “strong” hives surviving winter and “weak” hives not surviving winter), subspecies and geographical location. Pooling information and SRA accession numbers were described before (42). After sequencing, reads were quality controlled using Trimmomatic (49), version 0.38. Subsequently, *de novo* assemblies were made for the individual libraries using SPAdes (50), version 3.12.0, with kmer sizes 21, 33, 55 and 77 in the metagenomic mode. To remove redundancy, the resulting contigs larger than 500 bp were collapsed if they showed 97% nucleotide identity over at least 80% of the contig lengths, using ClusterGenomes (https://bitbucket.org/MAVERICLab/docker-clustergenomes). Putative eukaryotic viruses were identified using the BlastX method implemented in DIAMOND (43) version 0.9.22, using the ‘c 1’ and ‘sensitive’ flags, against the NR database (NCBI), downloaded on 30 september 2018. Taxonomical paths were parsed with the KtClassifyBLAST algorithm implemented in Kronatools (44). All contigs that fell under taxID ‘10239’ (Viruses) were included in the analysis. Contigs that could be annotated as bacteriophages (as described before (42)) were excluded. Coverage values per sample were obtained by mapping the reads per sample back to the viral dataset, using BWA-mem version 0.7.16a (51), filtering the obtained alignments for an identity of 97% over a coverage of 70% using BAMM (https://github.com/Ecogenomics/BamM). Coverage values were calculated by dividing the readcounts per contig by the contig length.

### SRA searches

The SRA database was searched by using the query ‘Hymenoptera + RNA’, and the resulting 5,246 fastQ files were retrieved by using the prefetch and fastq-dump tools implemented in the SRA toolkit (NCBI). The previously obtained viral dataset was used as an index and retrieved fastQ files were mapped back using BWA-mem (51), version 0.7.16a. Only samples that had a cumulative read count of at least 100,000 reads (1,331 samples) were included downstream. Samples were then *de novo* assembled using SKESA (46) and annotated and clustered as described above. Information on the included samples is provided in supplemental table 2.

### Phylogenetic analysis

Viral sequences were included based on an *ad hoc* determined length cut-off depending on the expected genome length of each virus (supplemental table S3). Reference sequences were included by using the retrieved viral sequences as query and performing a TBlastX search (52) with an e-value cutoff of 1E-10 against the nt database (NCBI), downloaded on 1 october 2019. For the Partiti-like, Tymo-like and Toti-like trees only Refseq sequences were included. Significant hits were also filtered on the abovementioned alignment length cut-off specific for a viral group (supplemental table 3). Next, proteins were predicted from both the queries and the significant hits, using prodigal (53), version 2.6.3. Predicted proteins were submitted to an all-to-all BlastP search, with an e-value cut-off of 1E-10. The output was then transformed into a network and the largest connected component was extracted using the networkx library (54) implemented in Python. Proteins within the largest connected component were subsequently aligned with MAFFT (55), version 7.313, using the L-INS-I setting and trimmed using trimAL (56), version 1.4.1, using the gappyout setting. Model selection was performed using Prottest (57), version 3.4.2. Bayesian phylogenetic analysis was performed using BEAST (45), version 1.10.4, using the predicted protein models (supplemental table 3) under a strict clock and constant population size prior. The respective analysis was ran until all the effective samples sizes were above 200, and maximum clade credibility trees were calculated using TreeAnnotator, implemented in the BEAST package. Final trees were plotted in R using the ggtree package (58).

### Network and contig sharing analysis

Networks were created from the retrieved viral sequence data by using TBlastX against the Refseq nt database, downloaded on 1 october 2019. The Refseq database was filtered by removing entries containing the keyword ‘phage’ (for bacteriophages) or ‘herpes’ in the header, and by removing sequences longer than 15000 nt and shorter than 500 nt. These cut-offs were implemented to reduce ‘noisy’ hits, where for example herpes polymerases have significant hits to other viral polymerases. The remaining sequences were clustered on 80% nucleotide identity over 80% of the length, by using CDhit, version 4.8.1 (59). The tBlastX search was performed with an E-value cutoff of 1E-10 and an alignment length cut-off of 300 positions, and was ran in two iterations to include reference sequences that only made the cut-off when aligning to other reference sequences. The resulting blast output was then converted into a minimized nested block network, using the graph-tool package (60), implemented in Python. Virus sharing over the eukaryotic families belonging to the Hymenoptera and within the Apidae was determined by using the coverage matrix. A viral sequence was assumed to originate from the taxonomical lineage of the sample of which the cluster representative (the longest contig inside a cluster) was derived, in order to determine directionality. A viral representative sequence was assumed to be present in a sample when the coverage was above 0.1. For the cumulative virus sharing, an additional threshold was imposed were at least 10% of the included samples of a specific taxonomical host lineage had to be positive before the viral sequence was assumed to be present within that lineage. Resulting networks were visualized in Cytoscape (61), version 3.7.1. Percentages of positive samples were calculated using the same relative count cutoff as mentioned before and the ASI and AMSI were calculated by taking the ratio of the fraction of positive samples for a specific bee virus within Apidae or *Apis Mellifera* samples, divided by the fraction of positive samples in other eukaryotic families or other *Apidae* species, respectively.

## QUANTIFICATION AND STATISTICAL ANALYSIS

PCoA analysis was performed in R (62) version 3.5.3, with the pcoa function implemented in the ‘ape’ library (63). Variance analysis and distance-based redundancy analysis was performed on the coverage matrix using Bray-Curtis distances, using the adonis test and capscale function implemented in vegan (64). Cumulative explanation power of the location (country of origin) and eukaryotic taxonomy (on family level) covariates was calculated using the ordiR2 function (vegan). The difference in absolute numbers of contigs was calculated using the Mann-Whitney U test implemented in scipy (65), in Python.

