## Supplemental Information for "Hymenoptera associated eukaryotic virome lacks host specificity"

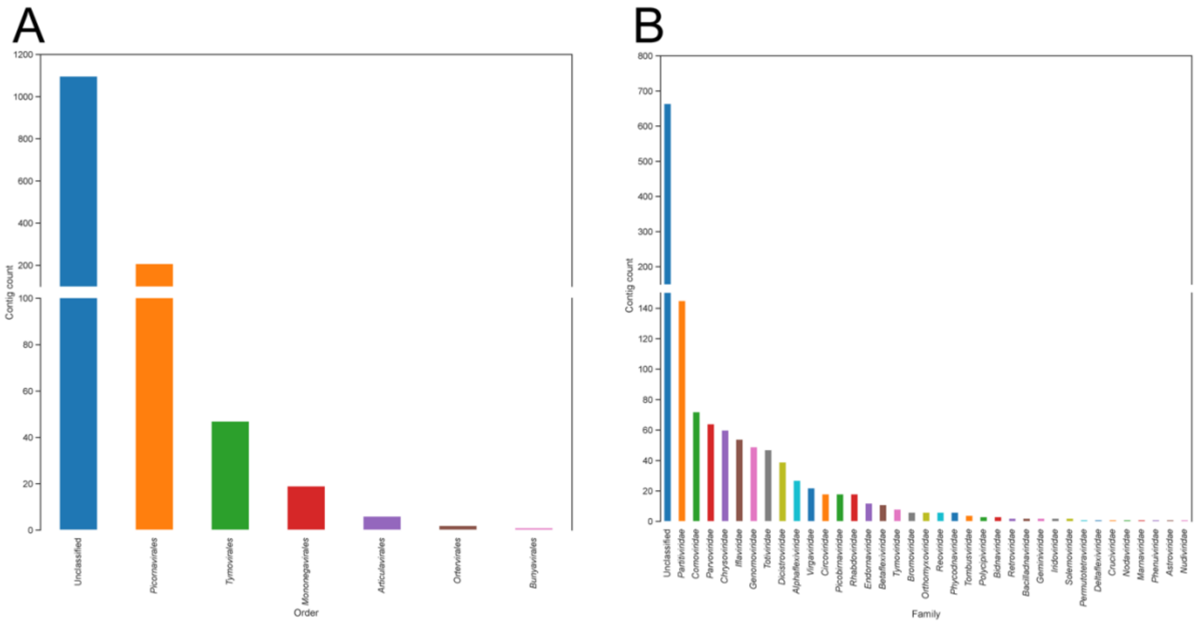


**Supplemental figure 1. Absolute contig count for the retrieved viral sequences from the Belgian samples**

Number of non-redundant contigs split per viral order (A) or viral family (B).

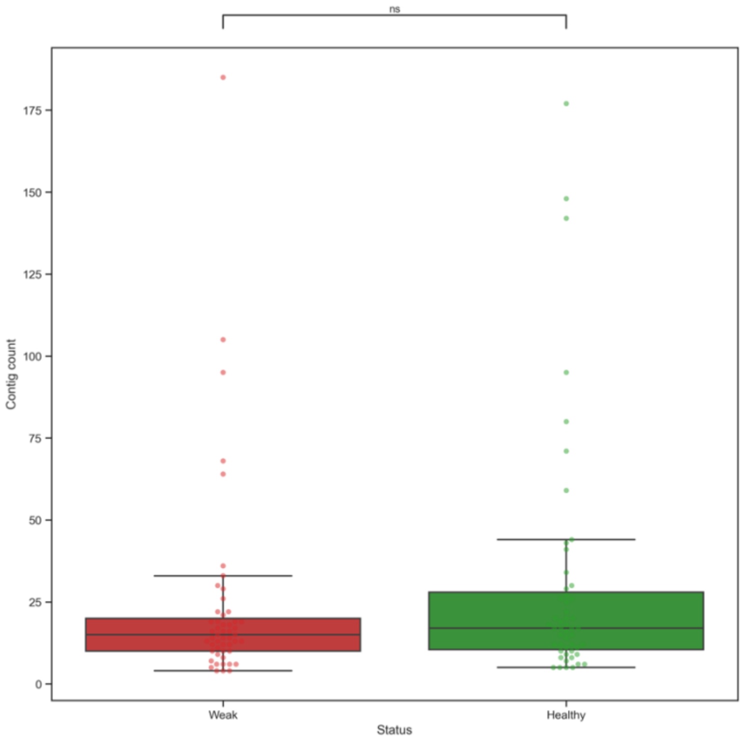


**Supplemental figure 2. No significant difference in number of viral contigs per sample status.**

Number of non-redundant contigs split per sample status (weak and healthy, n=51 for both groups). All data points are drawn. The box shows the three quartile values, and whiskers extend to 1.5 interquartile ranges. The horizontal bar above the plot indicates no significant difference (ns), tested with the two-sided Mann-Whitney U test (p-value = 0.30).

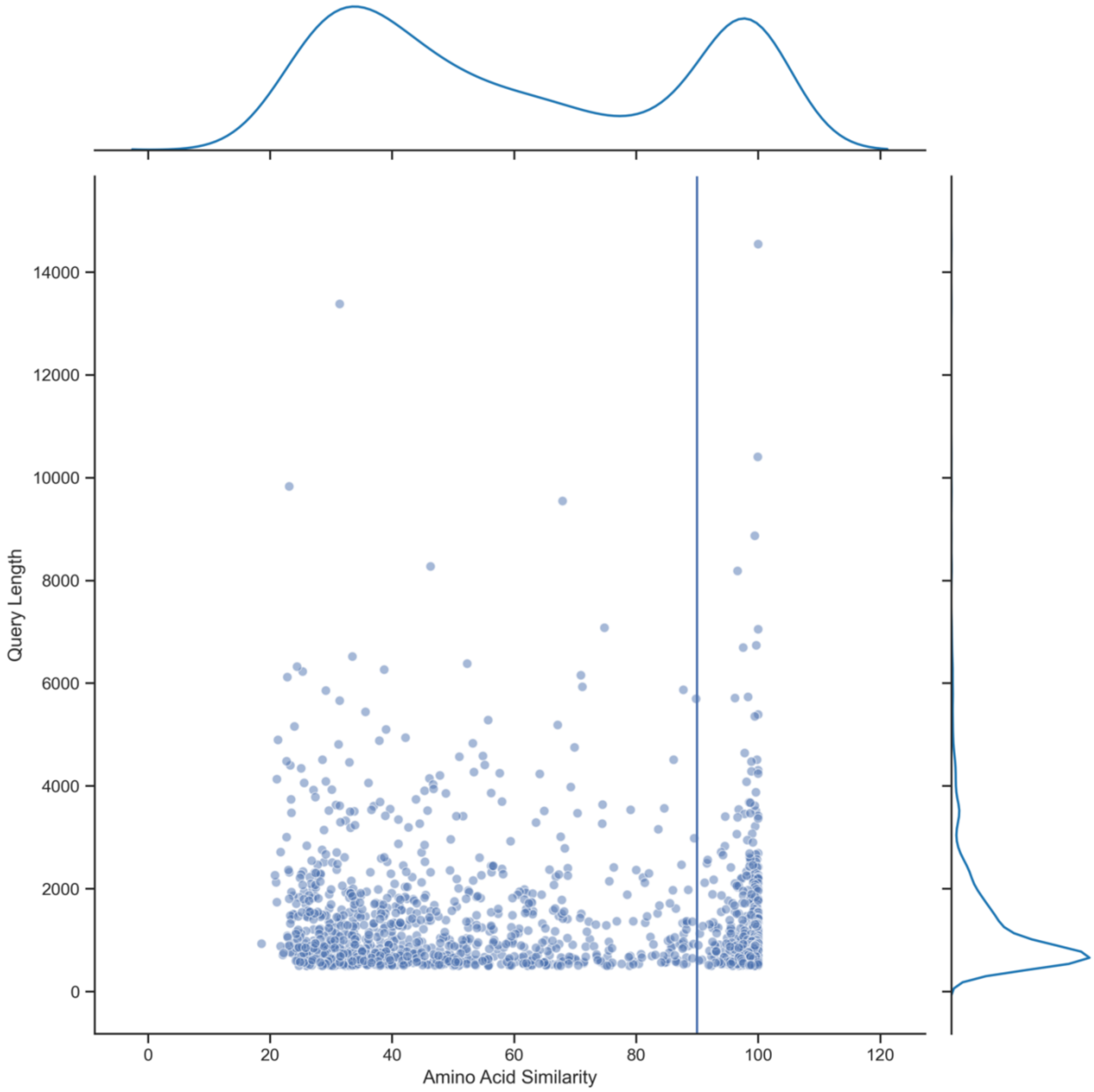


**Supplemental figure 3. Length versus similarity plots reveal the genetic divergent nature of most of the viral contigs from the Belgian samples.**
Scatterplot indicating contig length (in nucleotides) on the Y-axis, and the percentage of amino acid similarity (of the best hit) on the X-axis. Histograms in the marginal plots indicate densities. A vertical line is drawn on 90 amino acid similarity, the threshold for the 'known virus' group and 'unknown virus' group.

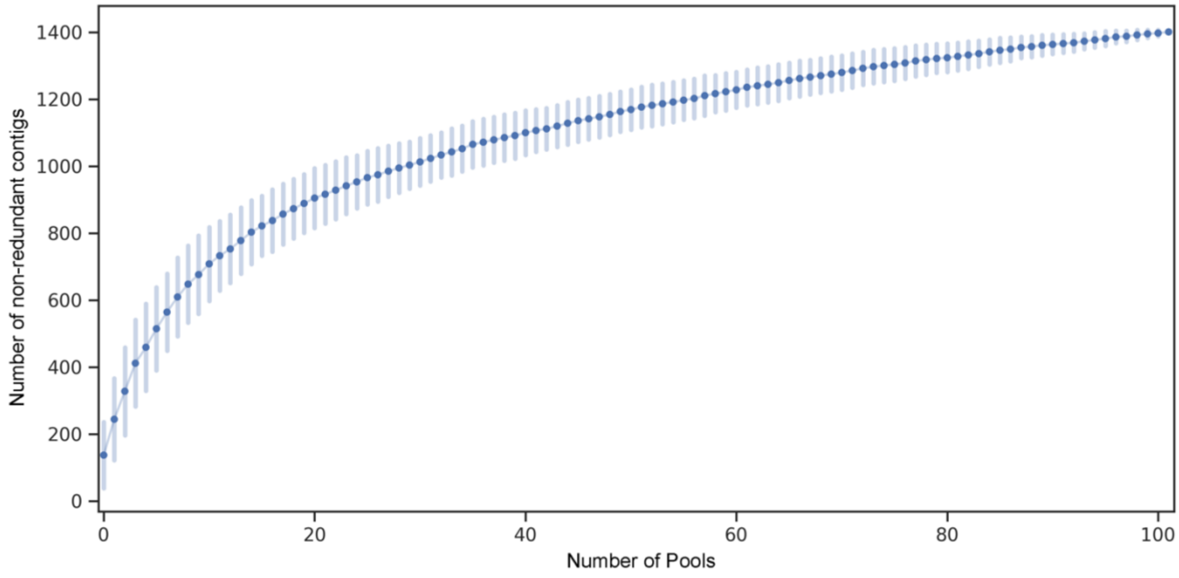


**Supplemental figure 4. Species accumulation curves reveal a plateau being reached.**
Species accumulation curves in function of the number of pools sequenced. Vertical lines indicate standard deviations based on 100 permutations.

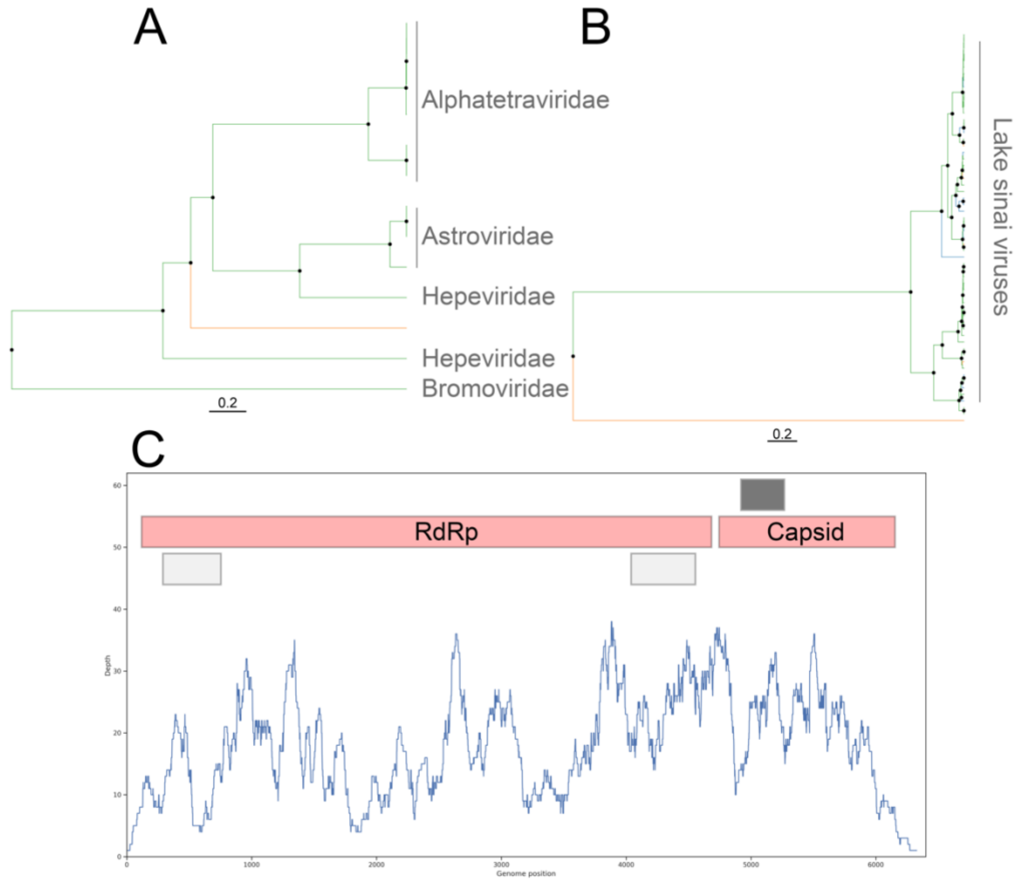


**Supplemental figure 5. Phylogenetic analysis and sequence depth profiling reveals the recombinant origin of LSV-Hepe-like virus.**
Panels A and B show the maximum clade credibility trees (based on amino acid alignments) for the putative polymerase region and the capsid region, respectively.

Reference sequences belonging to a classified viral family (left) or genus (right) are indicated in green. Orange branches indicate the LSV-Hepe-like virus. Nodes are labeled with a black circle when the posterior was higher than 0.9. Panel C shows the genome constellation of the putative LSV-Hepe-like virus. Red boxes indicate the putative polymerase (Non-structural) and capsid (Structural) genes, the grey boxes indicate unannotated predicted genes above 300 nucleotides in size. All putative ORFs are in sense orientation, with the exception of the two light gray ORFs. The curve shows the sequencing depth per nucleotide.

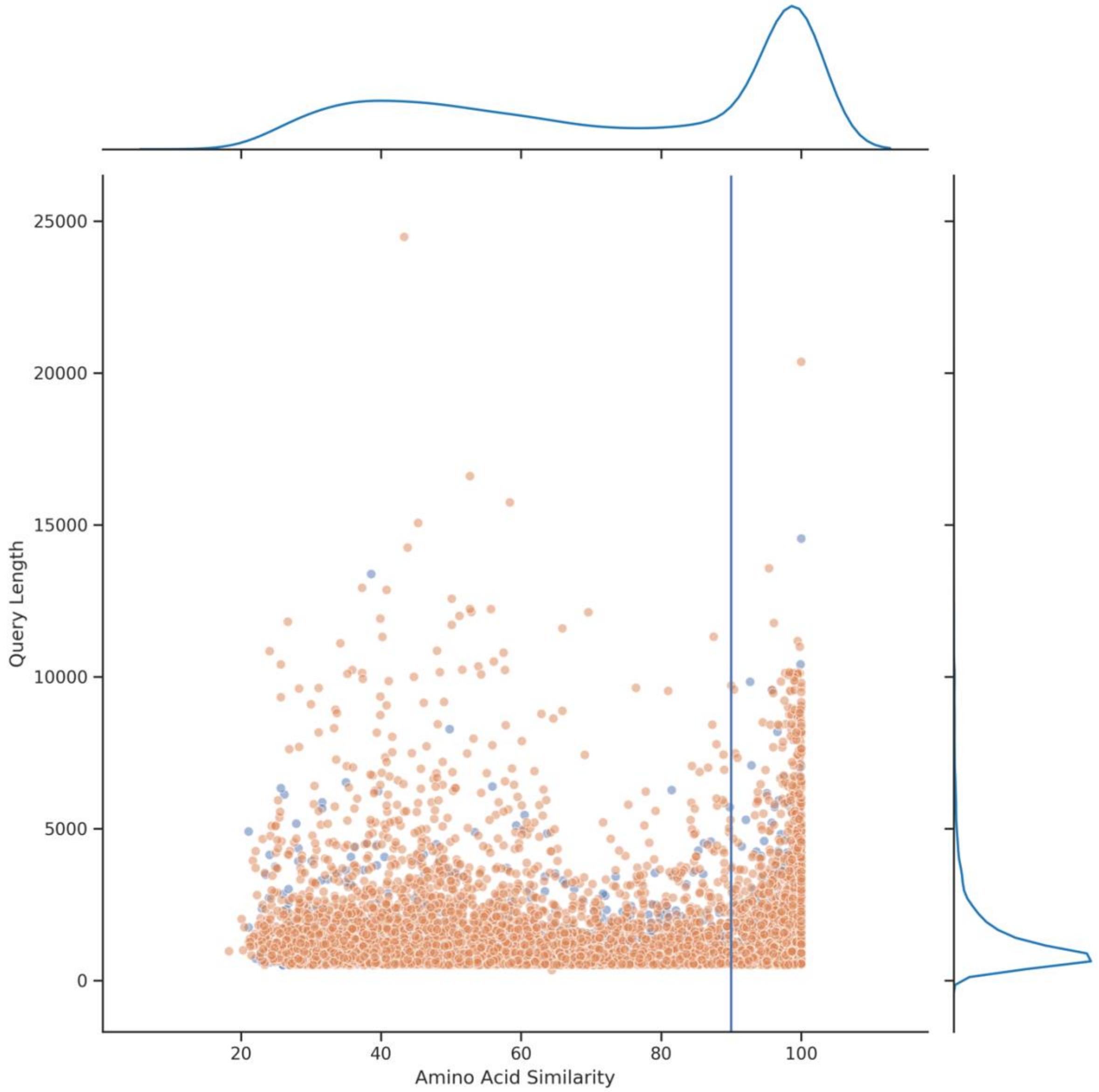


**Supplemental figure 6. Length versus Similarity plots reveal the divergence of most of the retrieved viral contigs from the Belgian samples and SRA searches.** Scatterplot indicating contig length (in nucleotides) on the Y-axis, and the percentage of amino acid similarity (of the best hit) on the X-axis. Marginal plots indicate KDE densities for both metrics. Orange dots reflect contigs identified in the SRA search, blue dots reflect contigs identified in the Belgian samples. A vertical line is drawn on 90 amino acid similarity, the threshold for the ‘known virus’ group and ‘unknown virus’ group.

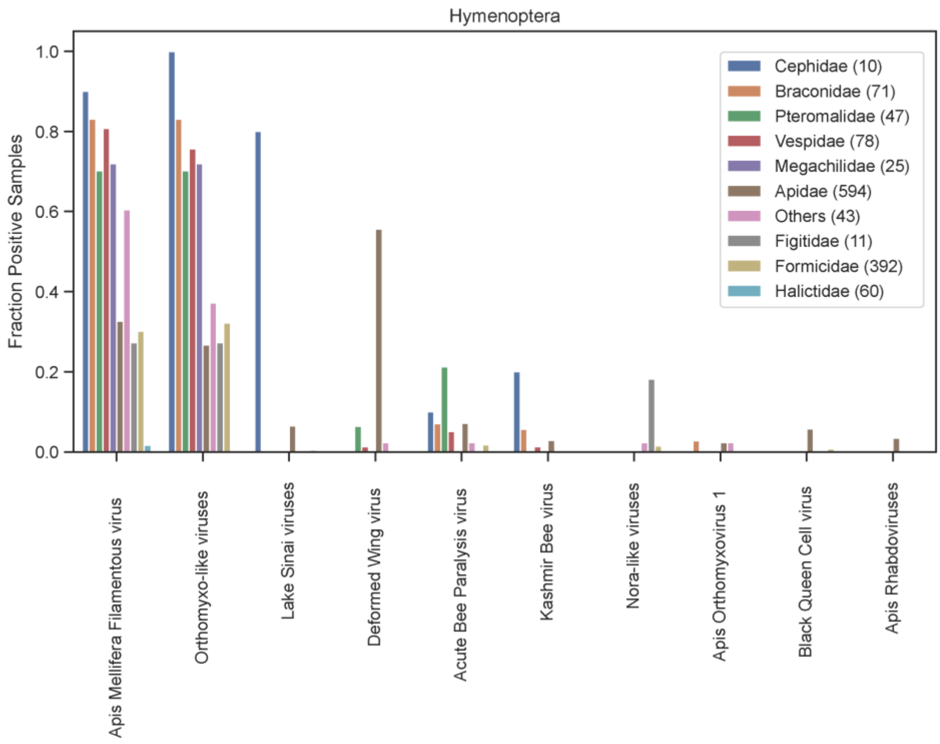


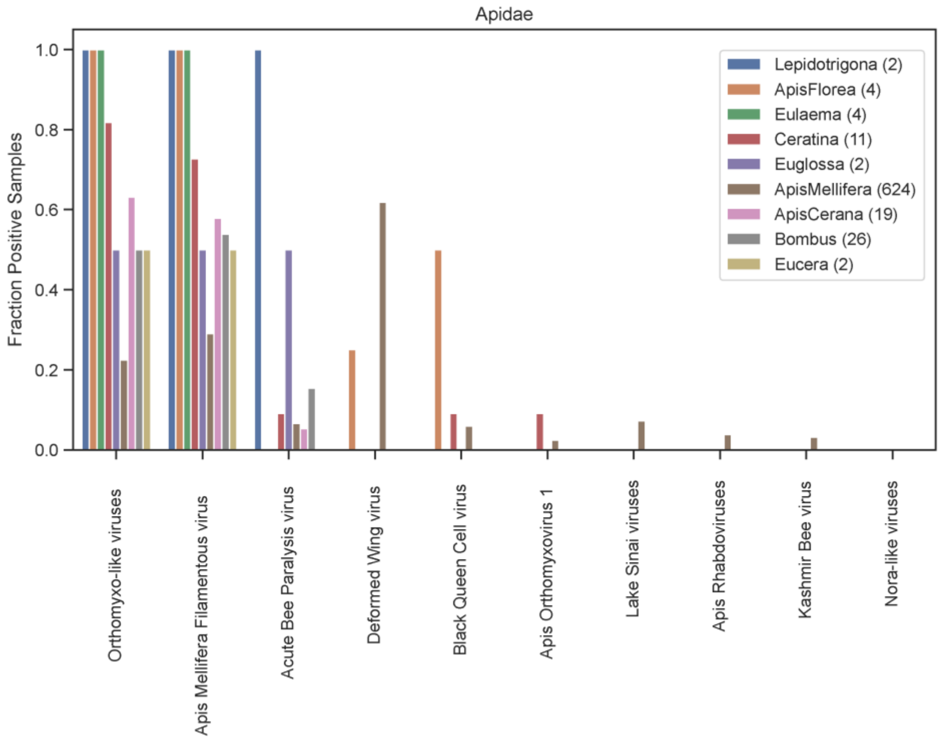


**Supplemental figure 7. Percentage of positive samples stratified per eukaryotic taxa imply that a substantial number of 'bee specific' viruses are not lineage restricted.**
Bar plots indicating the percentage of samples positive for a specific group of viruses. Different viral groups are indicated on the X-axis, the fraction of positive samples is denoted on the Y-axis. Barplots are colored according to eukaryotic lineage, either between hymenoptera families (top) or within the family Apidae (bottom). The sample size (number of datasets) per taxa is indicated in the legend.

**Supplemental table 1. Information on the viral contigs (taxonomy and GenBank accession number) derived from the Belgian samples and included in the phylogenies.**

**Supplemental table 2. Metadata for the public sequencing datasets that were included in the study. Additional columns on taxonomical information, geographical origin, date and reference study have been added and curated manually.**

**Supplemental table 3. Information on how the phylogenetic analysis was performed for the different viral groups. The threshold parameters for the TBlastX search and BlastP searches are indicated, as well as the protein model used, and the length of the chain in the BEAST runs.**
